# Neural Collision Detection: an open source library to study the three-dimensional interactions of neurons and other tree-like structures

**DOI:** 10.1101/2021.07.20.452894

**Authors:** Hagai Har-Gil, Yoav Jacobson, Alvar Prönneke, Jochen F. Staiger, Omri Tomer, Dan Halperin, Pablo Blinder

## Abstract

The analysis of neuronal structure and its relation to function has become a fundamental pillar in neuroscience since its earliest days, with the underlying premise that morphological properties can modulate neuronal computations. It is often the case that the rich three-dimensional structure of neurons is quantified by tools developed in other fields, such as graph theory and computational geometry; nevertheless, some of the more advanced tools developed in these fields have not yet been made accessible to the neuroscience community. Here we present Neural Collision Detection, a library providing high-level interfaces to collision-detection routines and alpha shape calculations, as well as statistical analysis and visualizations for 3D objects, with the aim to lower the entry gap for neuroscientists into these worlds. Our work here also demonstrates a variety of use cases for the library and exemplary analysis and visualizations that were carried out with it on real neuronal and vascular data.

## 1 Introduction

The unique morphology of types and sub-types of neurons in the mammalian brain has drawn researchers to extensively investigate their structure and its relation to function [1, 2]. Neurons have an elaborate network of connections between each other via neuron-to-neuron synapses, but they also continuously communicate with other cell types that surround them, including glial cells and those composing the neurovascular unit[3]. Thus, it is expected that the neuron’s geometric properties will also be affected, and could even be modulated, by the physical interactions with the cells on its surroundings [4, 5, 6], and vice versa.

This complex three-dimensional (3D) environment may be investigated from a variety of methodological approaches and angles. One such approach emphasizes geometrical interactions and uses *computational geometry* as its framework of reference [7, 8]. With this point of view neurons, for example, can be thought of as three dimensional rigid body graphs, with their soma being the parent node which branches out to different subgraphs that describe the axon and dendrites of that neuron. This structure is useful when seeking information which is closely tied to the morphological properties of the neuron and its surroundings, such as conductance along a branch, neuronal fiber distance between two spines or when modeling the interaction of a pair of neighbouring neurons. As a concrete example, Kashiwagi and colleagues [7] used computational-geometry-based methods to correlate evolving spine morphology with learning in an awake animal, and were able to find differences in the morphological structure between different strains of mice.

Questions such as the ones detailed above require designated *in silico* frameworks, either commercial or open-source ([9], as well as Neurolucida by MicroBrightFields, Inc., Colchester, VT, USA). A framework exposes common data structures and useful visualization tools and ultimately indirectly provides a set of questions that can be answered using its Application Programming Interface (API). However, any missing functionality might also mean that the framework’s users will be less inclined to ask questions which rely on that information, since it will necessitate them to either implement such functionality themselves or simply give up on such questions.

A prime example of such a missing feature is the gap between the single object level and macro-scale analysis [10, 11]. As noted, several existing frameworks have focused on the single cell level, letting their users analyze and visualize neurons, vasculature and other cell types in the brain with incredible precision. Other frameworks provide an API that assists with multicellular networks and modeling of cortical columns, providing answers to questions that relate to encoding and decoding of information in the brain [12, 13].

Motivated by the lack of computational environments that target cell-to-cell interactions and single-to-many relationships, we developed Neural Collision Detection (NCD) as a mean to bridge this gap. NCD’s main feature is a collision-detection engine capable of modeling the interaction of two 3D objects, for example a neuron with its surrounding vasculature, providing insight into their distance distribution, orientations and more. Furthermore, the library is able to batch process multiple two-object interactions, allowing to generate robust statistics. Finally, the library provides functionality to calculate the *alpha shape* spectrum [14] of the analyzed 3D objects. NCD is an open source Python and C++ package, which can be freely downloaded from our repository [15].

As an example of the novel questions NCD facilitates, we showcase here a spatially-implicit null hypotheses regarding the 3D arrangement of a single neuron within the vasculature that surrounds it. To the best of our knowledge, no other method has been used so far to generate the distribution of contacts between the two objects, which might point towards the more vascular-facing parts of neurons.

## 2 Materials and methods

### 2.1 Cell morphology data

Eighteen three-dimensional models of VIP-expressing GABAergic neurons, which form the majority of data from [2], were kindly received from the authors. These barrel-cortex neurons were imaged in the original study using structured illumination, with the raw data being post-processed by Neurolucida to form the final model of the cell. The authors of [2] also identified the exact cortical layer of each of the neurons using DAPI staining.

### 2.2 Vascular network data and neuronal positions from mouse somatosensory cortex

A cortical vascular network spanning 1053 by 987 by 1091 *μ*m^3^ from [16] was used. The network was segmented, skeletonized and its graph properties extracted using the software described in [17]. The data was converted to a triangle mesh[18], the format our collision-detection library requires, using custom Python and MATLAB scripts, which are freely available as part of our public library [15]. The positions of the neuronal *soma*(cell bodies), in the context of the vascular data, were also recorded and used in our computational pipeline to randomly position neurons for simulation purposes.

### 2.3 Collision detection library

At the heart of NCD sits its collision-detection algorithm, which relies upon the Flexible Collision Library (FCL) [19], a state-of-the-art software package for proximity queries. Below we detail the different configurations used in our FCL-based work (see also 7).

For its internal representation, FCL builds a model for each of the input objects. The objects are initially represented by triangle meshes, and for every object we construct an Oriented Bounding Box (OBB) [20] tree model out of these meshes (this is done once per model). An FCL model is a hierarchical tree of bounding boxes for sub-parts of the object, where the resolution increases deeper into the tree. As a result, the tree’s root stores a bounding box which envelopes the entire object, with each child node containing about half of the triangular mesh of the object until we reach a single triangle per leaf.

We use collision-detection to position a neuron from the aforementioned neuronal dataset within the complex vascular 3D network. The selected neuron is placed in the location of a randomly selected soma within the vascular data, in accordance to its original cortical layer. For each position, we seek to find the orientation that minimized (or results in zero) collisions between the neuronal dendritic and axonal arbor and the vasculature which is considered as a static object. The orientation of the neuron is set in a systematic fashion—a rotation in ℝ^3^ is defined by three angles, *α, β, γ*. When we rotate the neuron in our program, it is by *γ* degrees around the *z*-axis (parallel to the dorsal-ventral direction), then by *β* degrees around the *y*-axis and finally by *α* degrees around the *x*-axis. Biological constraints limit possible rotations such that |*α*| ≤ 5°, |*β*| ≤ 5°, while *γ* can be any angle. In addition, we work only with integer degrees, and use a resolution of 1°, which results in 43,560 possible orientations per position.

To test the collision of the object in a given position and orientation, FCL compares the trees of the objects and looks for two single triangles that intersect (see, e.g. [19] for details). This intersection is counted as a single collision and its position is marked as the position of one of the vertices of the mesh triangle which represents it. More details on this process are provided in section 3.1.4. Finally, NCD aggregates these points per orientation, position and object pairs to form a tabular database-like structure, where each row is a unique collision coordinate between two objects. This table can be parsed by accompanying scripts as part of our library.

### 2.4 Alpha shape calculation

A novel feature of NCD is its bindings to the Computational Geometry Algorithms Library (CGAL) [21], a state-of-the-art C++ computational geometry toolbox. NCD uses CGAL’s alpha shapes module to enhance the graph-structure of the 3D shape with information of its critical alpha value at every point. This added information allows the library users to correlate between the neuron’s structure and information regarding the concealment of each of its points, as explained in Section 3.3.

## 3 Results

### 3.1 Collision detection

A key feature of this library is its ability to efficiently and reliably detect collisions between two 3D objects. In addition, it is trivial to aggregate the collision-detection data from many pairs of objects and analyze the resulting trends. This collision process is composed of several key steps, outlined in Fig 1 and in the text below. The output of this step is a tabular data structure, with its rows being the collisions between each pair of objects, and the columns being properties of these collisions, such as their coordinates in space.

**Figure 1:**
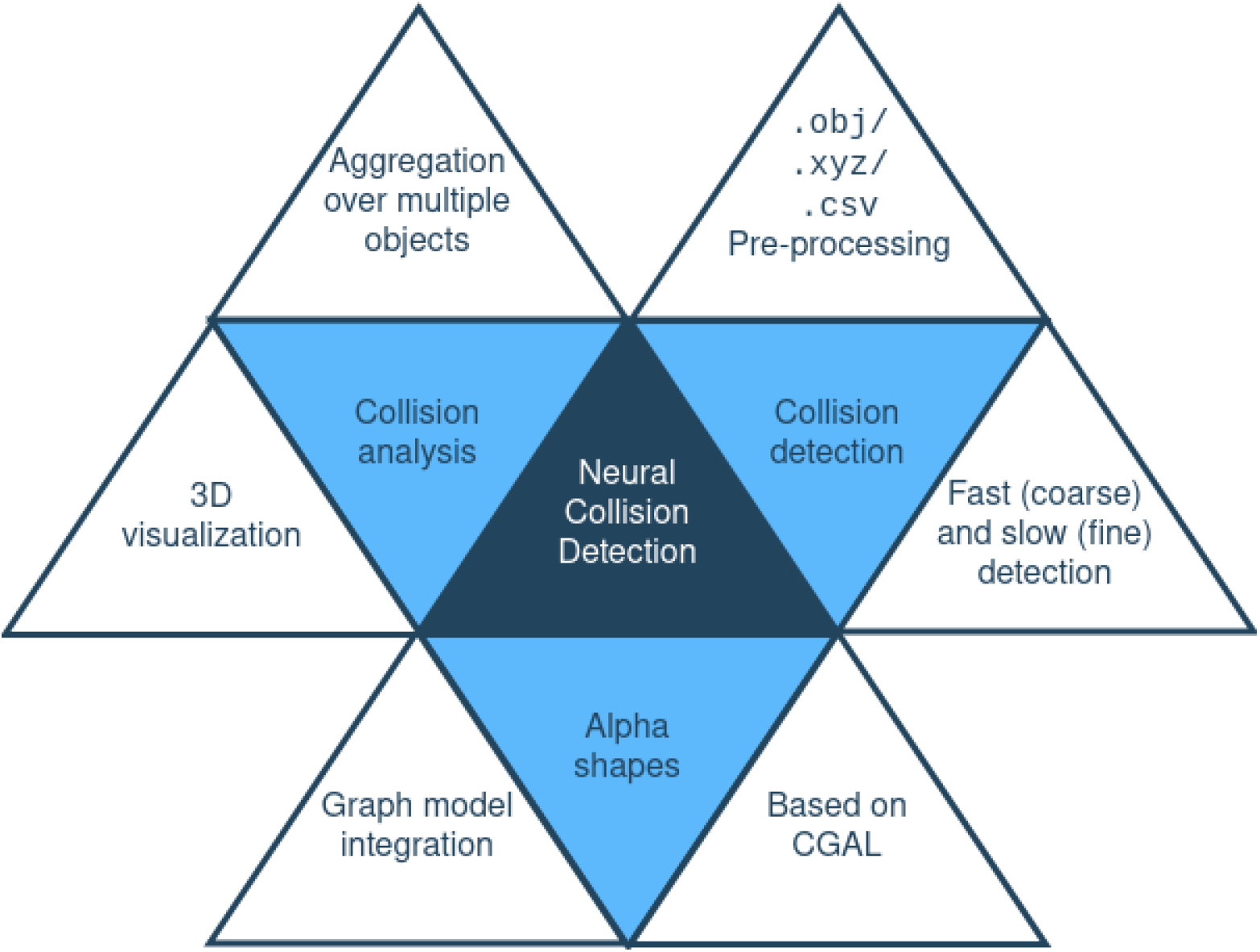
Package outline and its major capabilities. The tools this package provides can be divided into three key parts, highlighted in light blue. While these key parts can be used independently, their outputs are compatible with the inputs other parts are expecting. This means that, for example, visualizing the results of the collision-detection process may be done seamlessly due to this integration.

#### 3.1.1 Data pre-processing

NCD supports a wide variety of data formats that can be converted using existing functions to the format required by the collision-detection process itself. For example, point cloud-like tabular data can be converted to triangle meshes using built-in functionality, and vice-versa. By using multiple formats (and including conversion methods between them), NCD is able to support a range of output resolutions, which directly affect the tool’s performance, since a higher output resolution requires more processing power and computation steps.

#### 3.1.2 Collision detection process

Detecting collisions between two objects requires NCD to place them at some specific position and orientation, find the intersections and return either the coordinates of the intersections or their total number. NCD has a variety of parameters and modes that make this process very flexible and user friendly. For example, instead of re-running the software for each position and orientation one can supply a list of positions and orientations for one of the objects, while the other will stay put. Usually, to run it the user supplies the mesh representations of the two objects and a list of centers in which one of the objects should be placed at, while the other is stationary. NCD will then rotate the non-stationary object in multiple angles, as explained in Section 2.3 and record the number of collisions in each position or simply return their coordinates.

In this work we use a 3D vasculature model as the stationary input and a VIP interneuron as the dynamic one. The exact position of the VIP-expressing neuron’s soma is constrained to its corresponding cortical layer and to the set of soma-populated positions in the original dataset. In addition, the neuron must remain within the perimeter defined by the vasculature — we disallow any protrusions of neurites. These two 3D models are fed into NCD and the number of collisions per position and orientation is recorded for future filtering of collision-prone positions and orientations. At this stage, we discard the exact collision coordinates since we treat this step as a filteration step, meant to remove collision-prone positions and orientations. The exact collision coordinates will be computed later on using a more accurate algorithm, also encapsulated in NCD, that takes a longer time to run. Thus, per position, we are left only with the orientation having the least amount of collisions, and then these position-orientation pairs are sent to the aforementioned more precise functionality which returns the exact collision coordinates. Finally, the list of collision coordinates is passed downstream to be further aggregated and visualized, as seen in Fig 7 and explained in supplementary appendix 7.

#### 3.1.3 FCL-based coarse collision-detection procedure

To test for collision between two objects, NCD will fix one of them in place while translating and rotating the other according to user-defined parameters. The collision calculation is based on an OBB [20] model of the object, which is the preferred model when the collision process is heavily-based on rotations (as opposed to an Axis Aligned Bounding Box hierarchy [22]). In each orientation the triangles composing each model are compared and the number of pairwise intersections of the triangles are summed up.

This computationally-heavy part has been made more efficient by making sure that the objects’ model is created only once for the entire calculation and by running the computation for several positions in parallel, since they are independent of one another. The output of this step is the number of collisions per position-orientation pair.

#### 3.1.4 Refining collision coordinates

The FCL step explained above is a filtering step, designed to find the position-orientation pairs least likely to incur a collision. However the coordinates that the previous step returns are imprecise due to computational constraints. Namely, the data’s representation inside FCL can be a bit imprecise, and FCL’s output may not be “well-behaved”, as it may return points not residing on the actual objects. To refine this output we run a finer-grade computation which checks for collisions in a naive, “brute-force” method, resulting in a list of precise collision positions between the two objects, which in turn can be further post-processed by other parts of this library. This refinement step also supports different definitions of a collision by defining the threshold distance between the objects that counts as a collision.

### 3.2 Collision Analysis

The table resulting from the collision-detection step can be analyzed in a variety of ways. NCD supports a variety of post-processing steps, which use this table as their input. This provides composability of analysis functions, aiding the user to create complex post-processing pipelines from simple functions with a single purpose. For instance, a single function could aggregate all collisions occurring in some bounding box of interest, i.e. near the cell’s soma. This function will return its data in the same tabular format, with the irrelevant rows (collisions) filtered out. Then, another function could select only the rows with collisions that were closer than some fixed threshold values, and return another similar table. Each of these functions work independently on the same tabular format, but composing them together may spawn elaborate analysis pipelines.

#### 3.2.1 Neuron-as-graph with integrated collision data

It is common for neurons to be parsed into a graph structure to study their morphology [23]. This library offers such a tool with the added capability of integrating the collision probability of all nodes along the graph together with other graph-based properties. Example queries that can benefit from this extra information may include the correlation of synapses and/or buttons to the local collision probability, or the distribution of collisions along the dendrite, from the soma to its most remote parts.

#### 3.2.2 Visualization

The collision probability can be overlayed on top of a 3D visualization of the neuron, with or without the object that it collided with. These renderings may help researchers identify spatial patterns of the collision data as well as to further investigate more specific positions of the neuron that they might deem interesting, i.e. positions that caused collisions in specific areas of the models in question.

#### 3.2.3 Aggregation over multiple cells

Most analysis functions support a list of cells as inputs, instead of a single one. In some cases, such as visualization, this is a simple convenience, but in others it encourages running explicit comparisons between the neurons. For example, neurons from different layers might exhibit different correlations between the collision probabilities and their axons or dendrites.

#### 3.2.4 Exemplary visualization and analysis of collision data

Figs 2 and 3 show the capabilities of NCD to integrate collision and morphological data together via its underlying graph structure. We tested here for the collision of 16 neurons obtained from [2] with vascular data from [16] and analyzed the resulting distributions.

**Figure 2:**
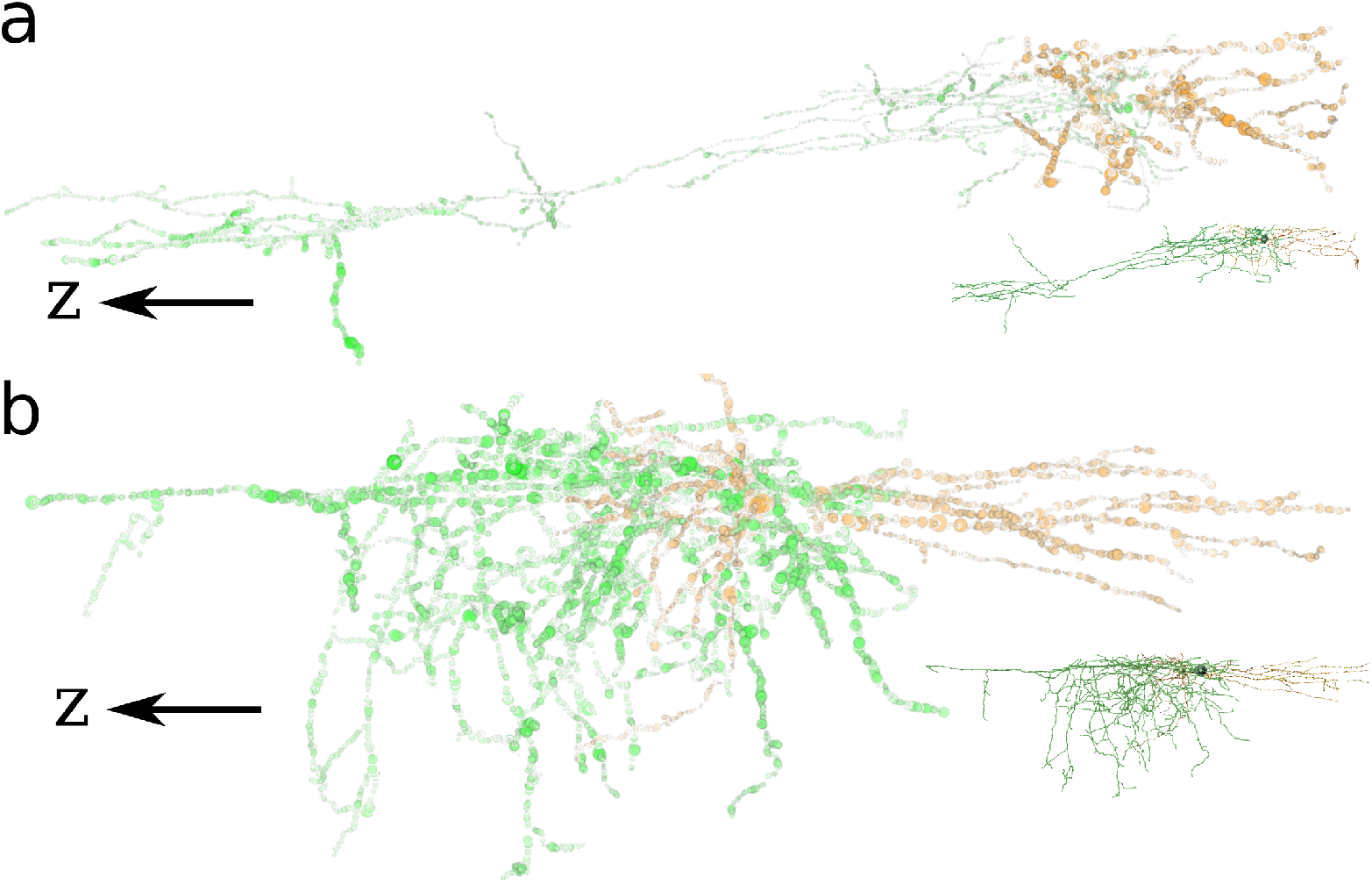
Exemplary visualizations of neuron collisions using NCD. 3D models of two representative cortical neurons from layer II\III (a) and layer V (b). The two distinct morphologies also generate a different collision distribution, as seen in the central part of neuron (a) which rarely collides with the surrounding vasculature. The *z* direction points toward the white matter (ventrally). The length of the arrow at (a) is 100 um, while the (b) arrow is 60 um. The napari application [24] was used for 3D renderings of the morphological structures and collision data. The insets show the neurons without collision data but with the same color coding, and with a dark sphere signaling the soma’s position.

**Figure 3:**
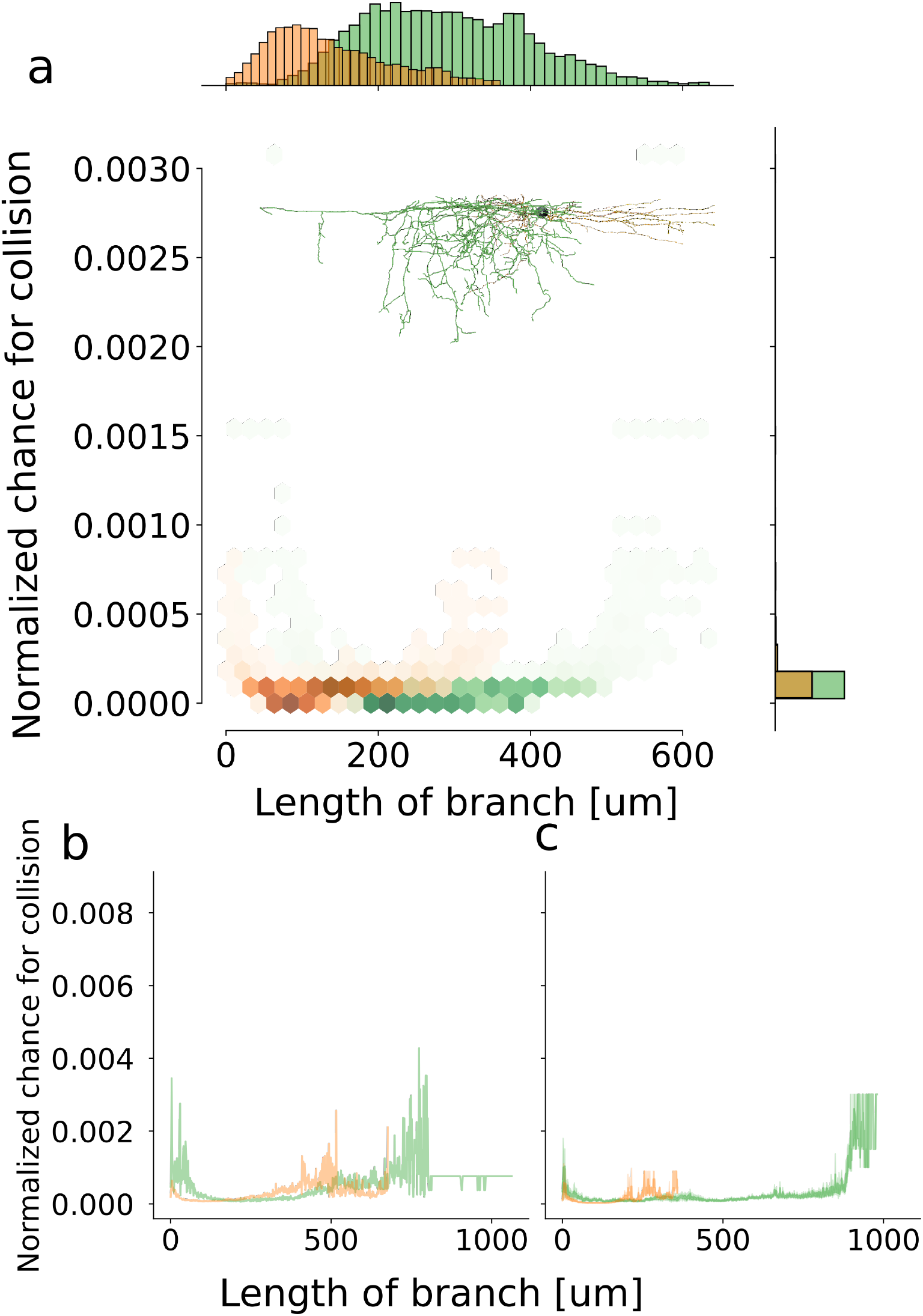
Collision probability as function of distance from the soma. (a) Example ‘hexbin’ density plot showcasing the collision distribution of dendritic (orange) and axonal population for the single neuron shown in the inset. (b-c) Layer II\III (b) collision distribution for the axonal (green) and dendritric population as opposed to layer V neurons (c). The dendrites of layer II\III neurons are generally more collision prone, while their axons show an increased collision probability at the middle and extreme parts.

First, in Fig 2, we see NCD-facilitated 3D renderings of neurons and their collision data. The two neurons shown, from layer II/III and from layer IV in the neocortex, were overlayed with their collision probabilities, with the size of the circle in correlation to the magnitude. The original 3D image of the neuron is also shown in the inset, and the colors code the axon (green) and dendrites (orange). The dark sphere in the insets marks the location of the cell’s body. These renderings may inform the viewer about disparities in collision probabilities between parts of the objects and can enable a more detailed analysis of the collision distribution.

Second, in Fig 3 we utilize the underlying graph structure to evaluate the distribution of collision probabilities for a single neuron and for groups of neurons. Panel (a) showcases the chance for a collision as we step further away from the soma for both axon (green) and dendrites (orange). The distributions are slightly shifted along the *x*-axis due to the different lengths of the axon and dendrites, but remain quite similar. The bottom part of the figure shows aggregations of that same data from neurons in layer II\III (b) and the neurons from other layers. This division highlights the dramatic change in morphology between the two subgroups, which is also visible in Fig 2, and how it affects the collision probability distribution.

The figures that showcase the collisions point to a few directions that library users can turn to when analyzing their own data. Viewing the results of the collision-detection step on top of the 3D reconstructions is achieved by integrating napari [24], an n-dimensional rendering library, with the generated data structures of NCD. This integration is quite seamless from the user’s persepective and can assist users to visually detect anomalies in their data. More quantitative strategies, as seen in Fig 3 are also possible, either at the single neuron level or at the population’s. Gathering and displaying these statistics easily is of utmost importance when trying to find trends in complex multidimensional data.

### 3.3 Exemplary analysis of alpha shapes

A compelling co-factor for a point on a neuron, especially in the context of collisions, is how hidden or concealed the point is in space, relative to other objects that occupy that same space. This parameter might be correlated to other geometric properties, and may explain some observed behavior between the two objects. Within NCD, as explained in Section 2.4, we offer the ability to compute alpha shapes [14] as a tool to quantify the topological structure of points, or nodes (in a graph sense), along the neuron.

In computational geometry, alpha shapes provide a way to capture the “shape” of a set of points, with the alpha parameter controlling how fine or crude the shape is. The alpha shape is a polytope that contains all of the points in a set, where some points from the set are its vertices. A point is a vertex of the alpha shape, if there exists a sphere (or a circle in 2D) of radius *α*, where the point lies on its boundary, and no point from the set is inside the sphere. In some sense, the more interior a point is, the lower the alpha parameters needs to be in order for the point to show up on the surface of the that alpha shape.

A point in a dataset has a alpha shape spectrum and critical alpha value. Its alpha shape spectrum is its categorizations over a range of different alpha shape values, whether the point is interior to the alpha shape or on its boundary. A critical alpha value is the minimal alpha radius that interiorizes the given point in the generated alpha shape. In other words, all larger alpha values treat that point as interior, and smaller values use this point as a part of their shell.

An example of two-dimensional analysis of alpha shapes for a small number of points is given in Fig 4. In the figure, the green zone marks the area covered by an alpha shape with some value — 0.7071 in the case of (a), and 0.7241 in the case of (b). These two values are critical alpha values since they represent a change in the coverage of the green area. With a lower alpha value (a) the alpha shape places the red dot near the center of the cross at its border, since the circles generated with the given radius are small enough to reach it. However, the larger circles that were used to form the area in (b) no longer touch the red dot, and are thus unable to use it in when setting the border. In this sense the inner red dot is more concealed that the rest of the black dots of the cross, besides the central one.

**Figure 4:**
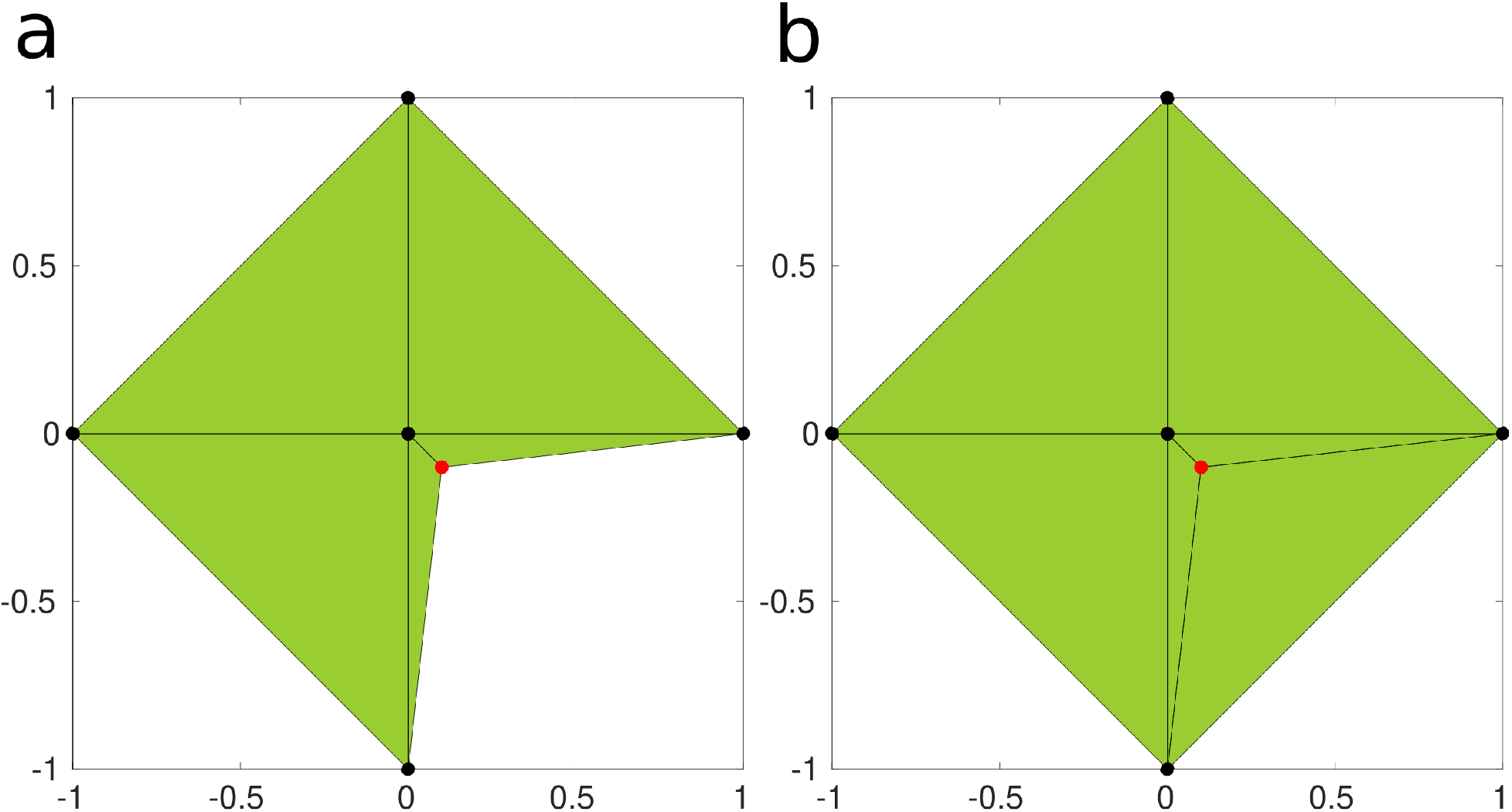
Two dimensional alpha shape analysis for six points in a plane. The red dot near the center of the cross, at (0.1, −0.1) simulates a concealed point and the green area connects between the borderline points of the alpha shape. (a) Calculation of the alpha spectrum reveals that alpha values between the critical values of 0.7071 and 0.724 will generate an alpha shape that is able to protrude deep enough into the cross so that it will include the concealed point in its border (alpha = 0.7071 is shown). (b) Alpha values larger than the critical value of 0.724 generate circles with a circumference that is unable to reach the inner point (alpha 0.7241 is shown). In other words, the inner point is concealed from that set of alpha shapes, namely it will not appear on the boundary of the alpha shapes for alpha greater than 0.724.

In our work, we pay less attention to the alpha shape itself, but rather focus on the points from the set that are vertices of the alpha shape for various alpha values, trying to capture how interior, or hidden, each point is. In the example above, the inner point has a low critical alpha value (0.724) while the points on the circumference will have a higher one, building the intuition that the red inner point is more hidden than the outer ones.

Our library is able to compute the alpha spectrum of each node on the neural graph and only record the largest alpha value of that spectrum, since it best represents the concealment (or “hiddenness” of that node. The alpha-value data is integrated into the neuron’s graph structure and is utilized by many analysis functions and scripts in the library.

As noted, NCD is able to calculate the alpha-shape spectrum for each node along the neuron using newly-generated Python bindings to the relevant components of the CGAL library. Furhter, NCD will also calculate the critical alpha value per point, as explained in Section 3.3.

Fig 5 shows the neurons from Fig 2 with the critical alpha radius overlayed onto their corresponding morphology. The insets show close-ups for specific areas in the neuron, which exhibit a significant difference between the alpha values of nearby axons and dendrites, suggesting that the morphological difference between these two nearby areas might bear biological significance. The largest radius in the image corresponds to the maximal allowed radius by the surrounding vasculature.

**Figure 5:**
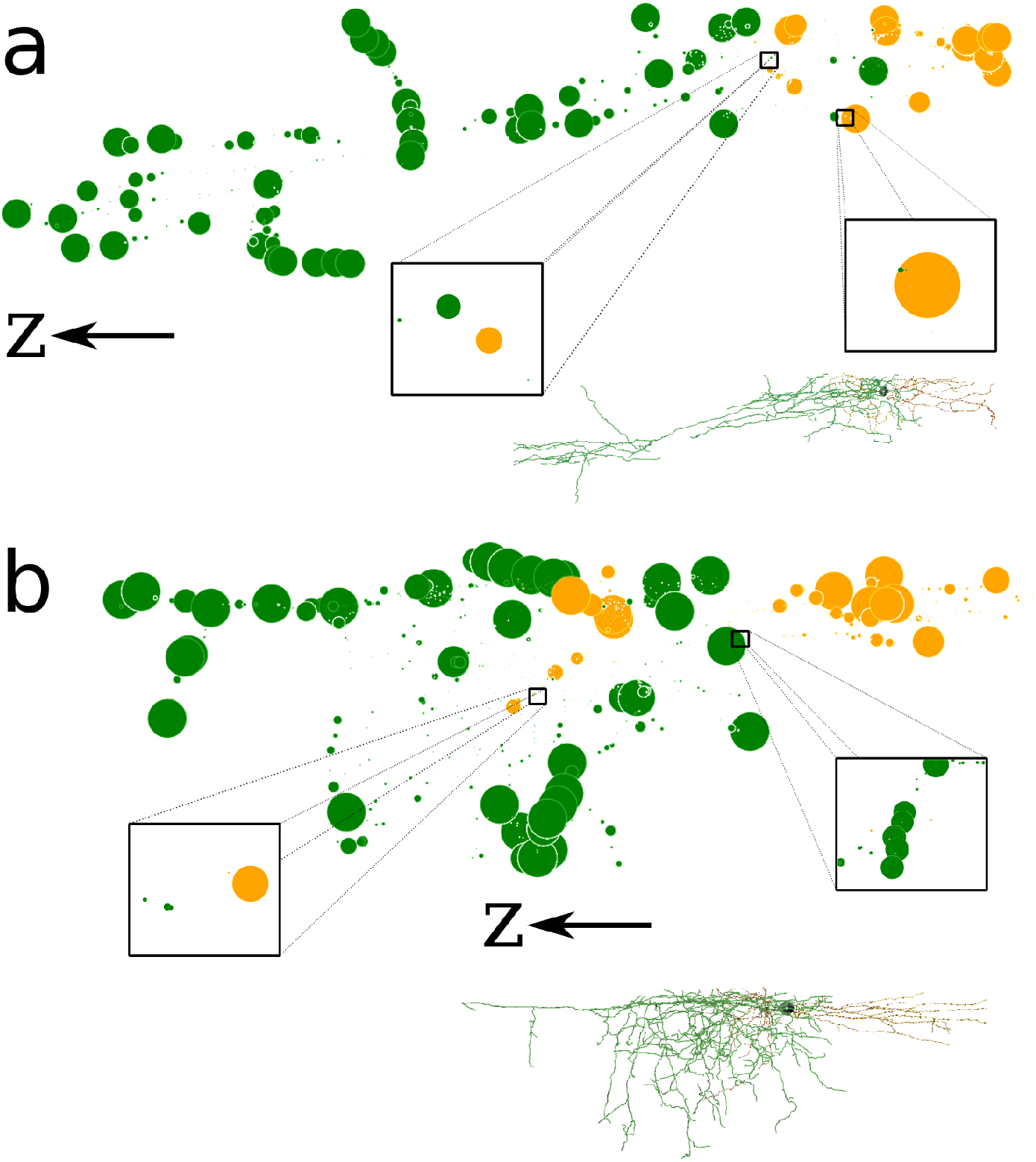
Exemplary 3D visualization of alpha values for the neurons shown in Fig 2. (a-b) The alpha values were integrated into the neuronal graph structure so that the size of the circle correlates to the alpha value of that point. The highlighted regions show how proximate regions of dendrites and/or axons can have very different corresponding alpha values. The *z* direction points toward the white matter (ventrally). The length of the arrow at (a) is 100 um, while the (b) arrow is 60 um. napari was used for 3D rendering.

On top of the visualization layer NCD also provides a large array of functions and scripts to analyze the alpha-shape critical values in relation to the other quantities that were introduced before. These include, for example, correlations between the alpha-shape values and the distance of each node to the soma. In addition, NCD can also compare the collision probability of each node to its alpha value and to other quantities (Fig 6).

**Figure 6:**
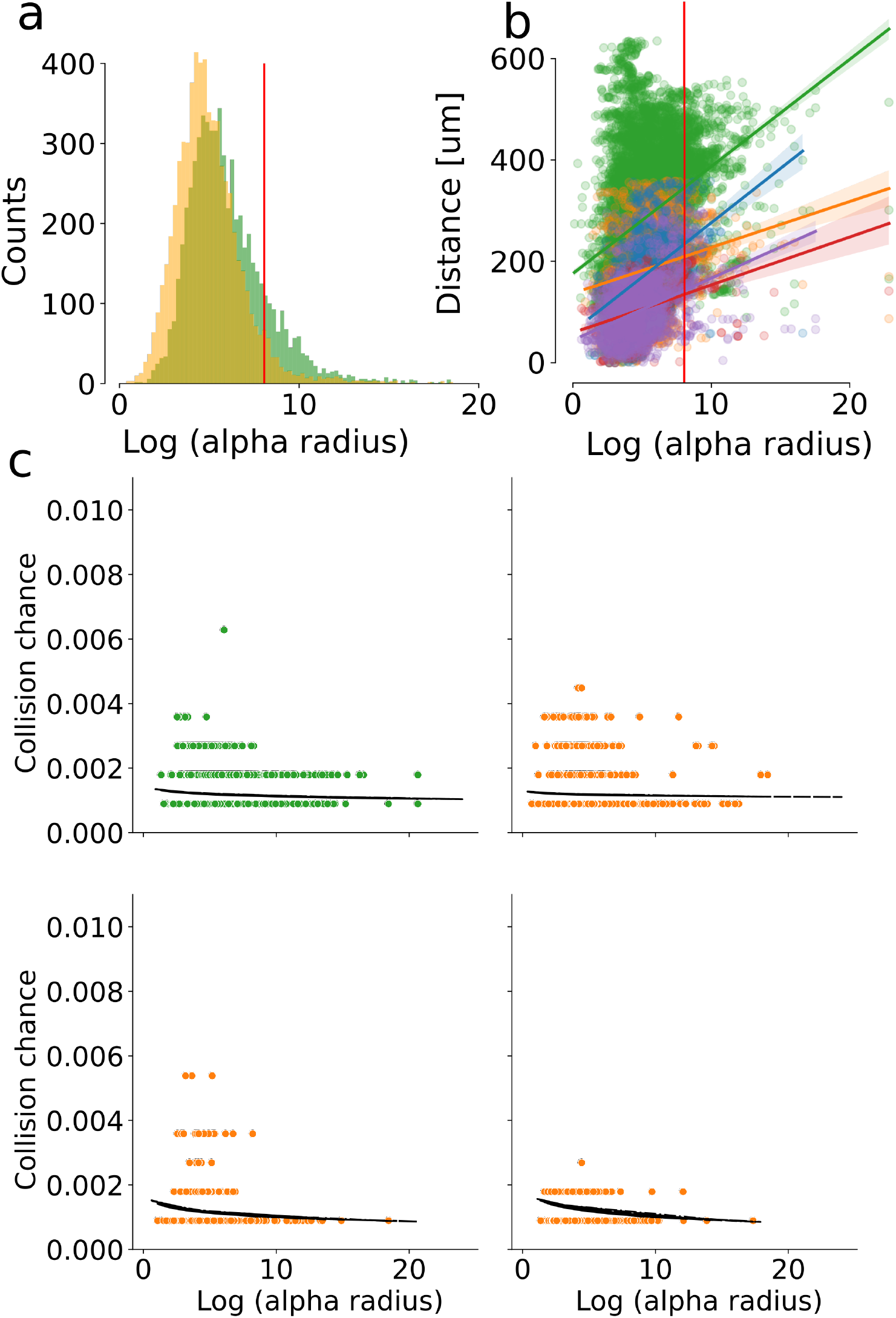
Alpha shape analysis examples using built-in NCD tools. (a) Critical alpha value distribution for a single neuron. The green histogram is for points on the axon while the orange depicts all dendrites. (b) The distribution of the topological distances from the soma as a function of the alpha values, with the respective trend lines. Green points are from the axon, while the rest of the points are colored by the specific dendritic tree they belong to. (c) Collision probability as a function of the critical alpha value for each point on the neuron, colored by neurite type. Data from the first two panels is from neuron (a) of Fig 2.

As seen in Fig 6, NCD facilitates a variety of analysis methods with its built-in functions. The neurons’ alpha-shape spectrum is pre-computed using novel CGAL bindings [15] and then this data can be further processed by functions in NCD’s `alpha_shapes` module. These methods include alpha-values distribution and correlation of the critical alpha values to other quantities such as topological distance and collision probability. In the figure the vertical red line points to the maximal vascular critical alpha value, which we use as the theoretical limit for the neurons’ alpha values once they are positioned inside the vasculature. Supplementary Fig 7 shows the alpha-value distribution of the vascular dataset that we used in this work.

## 4 Discussion

Our work showcases a new general software library for analysis aimed at solving neuroscientific questions using tools from computational geometry and robotics. This library, Neural Collision Detection (NCD), can be a one-stop shop for a variety of questions that deal with the interaction of two proximate units that populate the same tissue. Using its high performance “backends” in the form of the Flexible Collision Library and the Computational Geometry Algorithms Library, NCD is able to generate and integrate data and insights on the interactions of objects in 3D. Our tool also exposes computational pipelines that can provide insights from multiple pairs of such objects by aggregating the individual results and by providing helpful visualizations.

The NCD API exposes both high- and low-level functions to its users. Low-level functions grant users the ability to generate their own workflows and analysis scripts, while high-level ones are more opinionated in their approach to the data, which means that informative visualizations can be made with only a few lines of code. Together, these two classes of interfaces should appeal to a broad user base.

While extensive, NCD’s capabilities are not exhaustive. At present, the main limitation is its restriction on analyzing only two objects simultaneously. Batch analysis between multiple objects can be conducted, but each iteration will only include a pair of objects. Furthermore, NCD requires objects to be represented in specific formats. While we do supply converter functions, some formats are not covered in the currently-available scripts. Users are encouraged to contribute their converters to NCD itself or convert their data in a pre-processing step.

## 5 Future work

We believe that NCD can be used as a basic building block for more advanced processing pipelines. One such direction is real-time analysis of collision data, a requirement in fields such as robotics [22]. Since NCD is written in a high level language - Python - it can facilitate rapid prototyping of systems that require collision-detection but value speed of development over direct access to bare-metal constructs. Furthermore, NCD’s fast collision-detection mode is a convenient wrapper over FCL, which means that developers *pay* very little in terms of performance, but gain the ability to work with a simpler language than C++.

In addition, the faster collision-detection pipeline can be improved to provide even better performance. As an example, we can utilize the nature of the rotation process, and calculate collision for many positions at once [25] [26]. Improving the calculation-step speed may provide two somewhat conflicting benefits — it will prove NCD’s capabilities for real time work and may improve NCD’s collision-detection resolution in the first and fast phase.

It is also important to note that the slower collision-detection phase was developed due to the way we transformed the 3D neurons we worked with into their mesh representation. Due to the way the original data was captured, the transformations were not perfect, leaving discontinuities in the mesh representation. However, existing frameworks can usually streamline the correction of these artifacts in a user-friendly manner [27]. Moreover, we believe that better ground truths will account for better mesh representations, ones that might diffuse the need for the second collision-detection phase.

Lastly, the analysis steps shown in this work are more of a demonstration of future capabilities of the application. The complexity of the neuronal tree as well as the interactions between the neurons and blood vessels may cater a variety of research questions that are currently unanswered, like the relation between alpha values and collision probabilities in different neuronal populations, the mapping of collision sites on the vascular bed and more.

## Acknowledgments

The authors would like to thank Efi Fogel from the TAU CG Lab for his assistance in generating Python bindings for necessary parts of the C++-based CGAL. PB would also like to thank the support from the Israeli Science Foundation (1019/15) and the European Research Council (639416). Work by DH has been supported in part by the Israel Science Foundation (grant no. 1736/19), by NSF/US-Israel-BSF (grant no. 2019754), by the Israel Ministry of Science and Technology (grant no. 103129), by the Blavatnik Computer Science Research Fund, and by a grant from Yandex.

## 6 Author Contribution

HHG and YJ created the library, its API and bindings to FCL and CGAL and have written the manuscript. HHG had also conducted the computational simulations. AP and JS generated the neuronal data this work relies upon. OT created an early prototype of this work. DH and PB conceptualized the work, supervised its methodology, provided insights and assisted with the draft.

## 7 Supporting information

**Supplemental Figure 1:**
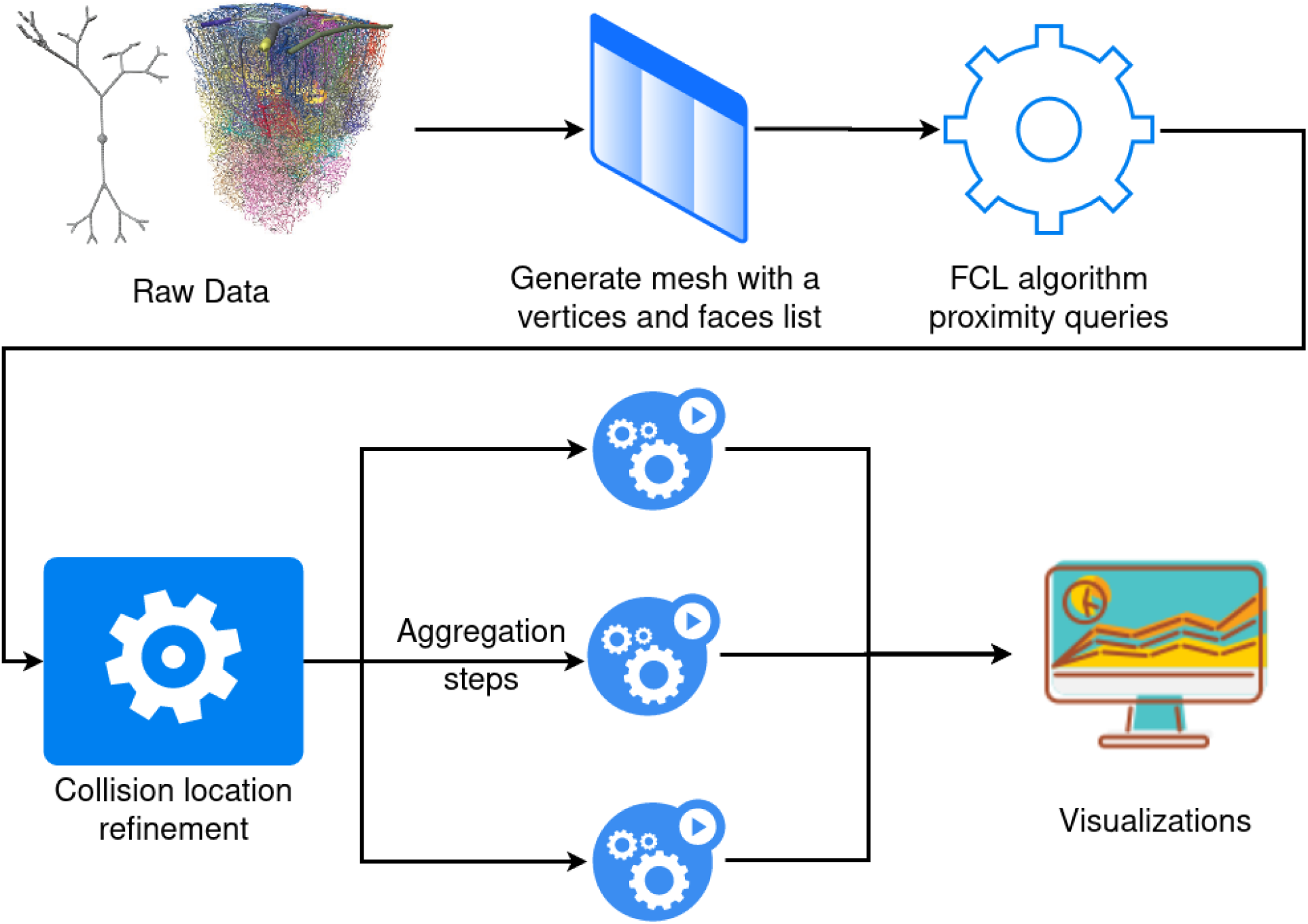
Explanation of the computational pipeline. The raw data is pre-processed into files which describe the 3D structure of the objects in the form of meshes. This type of data can be fed into FCL, which computes the crude positions of collisions between the neurons and vasculature. The calculation is first done in a coarse-grain manner due to performance reasons. After the approximate collision positions are detected, they are further refined by running a brute-force algorithm, which can also report the exact proximity of the neuron’s processes and the nearby vasculature. This brute-force method is slower to run, but it uses the previous crude step as a filtering method — exact collision point is found only for the points detected by the crude algorithm. Lastly, the properties of these interactions are aggregated and visualized, as described in this work.

**Supplemental Figure 2:**
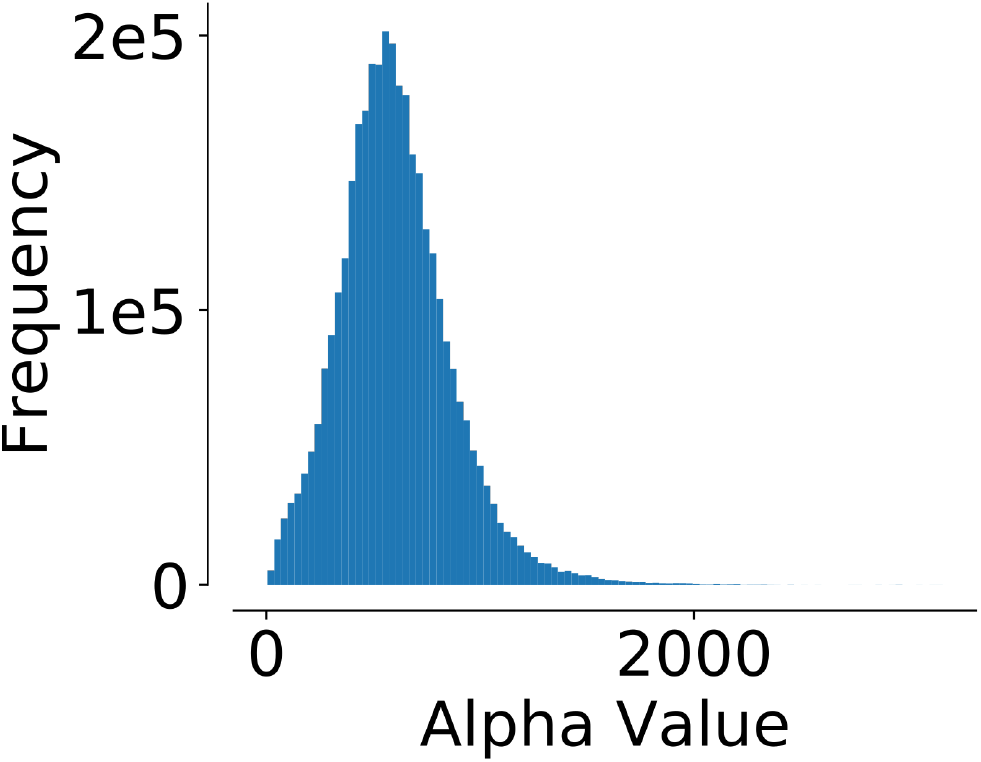
Critical alpha value for vasculature. The critical alpha-value for the each point on the vasculature model used in this work was calculated, and the distribution of these values is presented here. The maximal value of this histogram was used as a biological limit to the relevance of alpha-value calculation for neurons. For example, in cases where some points on the neuron exhibited a higher alpha-value than the maximal one calculated here, these values are considered biologically irrelevant since these so-called “exposed” areas will be concealed by the surrounding vasculature and by other cells in the parenchyma.

## References

[1] U.Valentin Nägerl, Nicola Eberhorn, Sidney B. Cambridge, and Tobias Bonhoeffer. Bidirectional activity-dependent morphological plasticity in hippocampal neurons. Neuron, 44(5):759 – 767, 2004.

[2] Alvar Prönneke, Bianca Scheuer, Robin J. Wagener, Martin Möck, Mirko Witte, and Jochen F. Staiger. Characterizing VIP Neurons in the Barrel Cortex of VIPcre/tdTomato Mice Reveals Layer-Specific Differences. Cerebral Cortex, 25(12):4854–4868, 09 2015.

[3] Bruno Cauli, Xin-Kang Tong, Armelle Rancillac, Nella Serluca, Bertrand Lambolez, Jean Rossier, and Edith Hamel. Cortical gaba interneurons in neurovascular coupling: Relays for subcortical vasoactive pathways. Journal of Neuroscience, 24(41):8940–8949, 2004.

[4] Netanel Ofer and Orit Shefi. Axonal geometry as a tool for modulating firing patterns. Applied Mathematical Modelling, 40(4):3175 – 3184, 2016.

[5] Joseph Bastian and Jerry Nguyenkim. Dendritic modulation of burst-like firing in sensory neurons. Journal of Neurophysiology, 85(1):10–22, 2001. PMID: 11152701.

[6] Verónica C. Piatti, M. Georgina Davies-Sala, M. Soledad Espósito, Lucas A. Mongiat, Mariela F. Trinchero, and Alejandro F. Schinder. The timing for neuronal maturation in the adult hippocampus is modulated by local network activity. Journal of Neuroscience, 31(21):7715–7728, 2011.

[7] Yutaro Kashiwagi, Takahito Higashi, Kazuki Obashi, Yuka Sato, Noboru H. Komiyama, Seth G. N. Grant, and Shigeo Okabe. Computational geometry analysis of dendritic spines by structured illumination microscopy. Nature Communications, 10(1):1285, Mar 2019.

[8] Sharon A. Swanger, Xiaodi Yao, Christina Gross, and Gary J. Bassell. Automated 4d analysis of dendritic spine morphology: applications to stimulus-induced spine remodeling and pharmacological rescue in a disease model. Molecular Brain, 4(1):38, Oct 2011.

[9] Hermann Cuntz, Friedrich Forstner, Alexander Borst, and Michael Häusser. One rule to grow them all: a general theory of neuronal branching and its practical application. PLoS Comput Biol, 6(8):e1000877, 2010.

[10] Alexander Shakeel Bates, James D Manton, Sridhar R Jagannathan, Marta Costa, Philipp Schlegel, Torsten Rohlfing, and Gregory SXE Jefferis. The natverse, a versatile toolbox for combining and analysing neuroanatomical data. Elife, 9:e53350, 2020.

[11] Ju Lu. Neuronal tracing for connectomic studies. Neuroinformatics, 9(2-3):159–166, 2011.

[12] J. Dyhrfjeld-Johnsen, J. Maier, D. Schubert, J. Staiger, H.J. Luhmann, K.E. Stephan, and R. Kötter. Cocodat: a database system for organizing and selecting quantitative data on single neurons and neuronal microcircuitry. Journal of Neuroscience Methods, 141(2):291–308, 2005.

[13] Blue Brain Project. Morphio. https://github.com/BlueBrain/MorphIO/, 2021.

[14] Herbert Edelsbrunner and Ernst P Mücke. Three-dimensional alpha shapes. ACM Transactions on Graphics (TOG), 13(1):43–72, 1994.

[15] Dan Halperin Yoav Jacobson, Hagai Har-Gil and Pablo Blinder. Neural collision detection. https://github.com/PBLab/neural_collision_detection, 2021.

[16] Pablo Blinder, Philbert S Tsai, John P Kaufhold, Per M Knutsen, Harry Suhl, and David Kleinfeld. The cortical angiome: an interconnected vascular network with noncolumnar patterns of blood flow. Nature neuroscience, 16(7):889–97, jul 2013.

[17] Philbert S Tsai, John P Kaufhold, Pablo Blinder, Beth Friedman, Patrick J Drew, Harvey J Karten, Patrick D Lyden, and David Kleinfeld. Correlations of neuronal and microvascular densities in murine cortex revealed by direct counting and colocalization of nuclei and vessels. Journal of Neuroscience, 29(46):14553–14570, 2009.

[18] Mario Botsch, Leif Kobbelt, Mark Pauly, Pierre Alliez, and Bruno Lévy. Polygon Mesh Processing. A K Peters, 2010.

[19] J. Pan, S. Chitta, and D. Manocha. FCL: A general purpose library for collision and proximity queries (commit hash 3f5963). In 2012 IEEE International Conference on Robotics and Automation, pages 3859–3866, May 2012.

[20] Stefan Gottschalk, Ming C. Lin, and Dinesh Manocha. Obbtree: A hierarchical structure for rapid interference detection. In Proceedings of the 23rd Annual Conference on Computer Graphics and Interactive Techniques, SIGGRAPH 1996, New Orleans, LA, USA, August 4-9, 1996, pages 171–180, 1996.

[21] The CGAL Project. CGAL User and Reference Manual. CGAL Editorial Board, 5.1 edition, 2020.

[22] Ming C. Lin, Dinesh Manocha, and Young J. Kim. Collision and proximity queries. In Handbook of Discrete and Computational Geometry, chapter 39, pages 1029–1056. Chapman & Hall/CRC, 2018.

[23] Paulo Aguiar, Mafalda Sousa, and Peter Szucs. Versatile morphometric analysis and visualization of the three-dimensional structure of neurons. Neuroinformatics, 11(4):393–403, 2013.

[24] Nicholas Sofroniew, Talley Lambert, Kira Evans, Juan Nunez-Iglesias, Kevin Yamauchi, Ahmet Can Solak, Philip Winston, Grzegorz Bokota, ziyangczi, Genevieve Buckley, Tony Tung, Draga Doncila Pop, Hector, Jeremy Freeman, Matthias Bussonnier, Peter Boone, Loic Royer, Hagai Har-Gil, Alan R Lowe, Mark Kittisopikul, Shannon Axelrod, Ariel Rokem, Bryant, Christoph Gohlke, Justin Kiggins, Mars Huang, Pranathi Vemuri, Reece Dunham, Trevor Manz, and Volker Hilsenstein. napari/napari: 0.4.0, October 2020.

[25] Stephen Cameron. Collision detection by four-dimensional intersection testing. IEEE Transactions on Robotics and Automation, 6(3):291–302, 1990.

[26] Karim Abdel-Malek, Jingzhou Yang, Denis Blackmore, and Ken Joy. Swept volumes: fundation, perspectives, and applications. International Journal of Shape Modeling, 12(01):87–127, 2006.

[27] Marwan Abdellah, Juan Hernando, Stefan Eilemann, Samuel Lapere, Nicolas Antille, Henry Markram, and Felix Schürmann. NeuroMorphoVis: a collaborative framework for analysis and visualization of neuronal morphology skeletons reconstructed from microscopy stacks. Bioinformatics, 34(13):i574–i582, 06 2018.

